# A data-based guide to the North American ecology faculty job market

**DOI:** 10.1101/751867

**Authors:** Jeremy Fox

## Abstract

Every year for three years (2016 to 2018), I tried to identify every single person hired as a tenure track prof in ecology or an allied field (e.g., fish & wildlife) in N. America. I identified a total of 566 hires. I used public sources to compile various data on the new hires and the institutions that hired them (e.g., number of publications, teaching experience, hiring institution Carnegie class). I also compiled data provided by anonymous ecology faculty job seekers on ecoevojobs.net (e.g., number of positions applied for, number of publications, numbers of interviews and offers). And I polled readers of the Dynamic Ecology blog to get information about applicant and search committee behavior (e.g., regarding customization of applications to the hiring institution). These data address some widespread anxieties and misunderstandings about the ecology faculty job market, and also speak to gender diversity and equity in recent ecology faculty hiring. They complement, and in some cases improve on, other sources of information, such as anecdotal personal experiences.

## Introduction

Academia is among the common career paths for Ph.D.-holding ecologists in North America (Hampton and Labou 2017).^1^ Some additional fraction of Ph.D.-holding ecologists presumably seek a faculty position at some point.

Ecology faculty job seekers want information about the ecology faculty job market. Indeed, there’s a widespread perception that they already have it. That because academia was traditionally seen as the “default” career path for ecology Ph.D.s, good information about that career path is widely and easily available. But in fact, good systemic information either doesn’t exist, or exists only for fields that are unlike ecology in important ways. So all ecology faculty job seekers have to go on are their own and others’ anecdotal personal experiences, and advice from others that itself is based on anecdotal experiences. Anecdotes, and advice based on them, can be useful sources of information. But they also can be unrepresentative or otherwise misleading, like any small and non-random sample. The competitiveness of the ecology faculty job market combined with the lack of good systemic information creates a fertile breeding ground for rumors and misplaced anxiety.

Both ecology faculty job holders and job seekers care about diversity and equity in faculty hiring. Overt and subconscious discrimination, and systemic forces that shape individual decision-making, can prevent the best people from applying, and can prevent the best people from being hired if they apply. Further, academic departments, and the colleges and universities comprised of them, are institutional wholes that are greater than the sum of their parts. Those institutional wholes are best able to teach and inspire the full range of students who come through their doors, and best able to pursue new knowledge, if they’re comprised of diverse, complementary mixtures of people. However, the most oft-cited data on faculty diversity concern the diversity of current faculty. Such data are informative about the ongoing legacy of past inequitable hiring. But they are uninformative about diversity and equity in current hiring because most current faculty were hired a decade or more ago. Available data on faculty diversity also are aggregated into broad fields such as biology, and so are uninformative about ecology specifically. We need data on recent faculty hiring in ecology to evaluate whether current faculty hiring in ecology is diverse and equitable at a systemic level, and to focus solutions on problems that exist in ecology rather than on problems specific to other fields.

Here I present a statistical profile of the North American ecology tenure-track faculty job market. I provide systemic data on the available positions, hiring institutions, applicants, and new hires.

## Data sources

Data about positions and applicants came from ecoevojobs.net. It is a crowdsourced spreadsheet which links to and summarizes advertised faculty positions in ecology and allied fields. A new spreadsheet is produced annually for each “job year”, starting in the early summer of the calendar year. The listing of tenure-track North American assistant professor positions in ecology and allied fields is nearly comprehensive. Unless otherwise noted, all my data come from the 2015-16, 2016-17, and 2017-18 spreadsheets. Ecoevojobs.net also allows anonymous ecology faculty job seekers to publicly share information about themselves and the positions for which they applied, including but not limited to year of Ph.D., current position, gender, race/ethnicity, number of publications (first-authored and other), number of applications submitted, number of interviews received (phone/skype and on campus), and number of offers received.

To obtain data on new hires, each year from 2016 to 2018, in the summer and fall following the end of the “job year”, I attempted to identify everyone hired into a North American tenure-track assistant professor position in ecology or an allied field. Starting from the ecoevojobs.net position list of tenure-track positions, I first culled any position that seemed unlikely to be filled by an ecologist, based on the job title (e.g., genomics, cell biology, anatomy & physiology). The retained positions were in areas such as ecology, fisheries, wildlife, natural resources, forestry, conservation, botany, zoology, entomology, and biology. I also culled any job that couldn’t be filled at the assistant professor level (e.g., ads for department chairs). Each year, that left me with approximately 300 tenure-track assistant professor positions that could potentially be filled by ecologists. I attempted to determine who, if anyone, was hired into those jobs by looking at department websites, emailing colleagues and department chairs, posting requests for information on the Dynamic Ecology blog and the Ecolog-L listserv, and searching the web and social media. I included any spousal hires I was told about.

For every position I was able to identify, I recorded if it was filled by an ecologist at the assistant professor level, filled in some other way (e.g., filled by a non-ecologist, filled at the associate level), or not filled. Who to count as an “ecologist” (as opposed to, say, an evolutionary biologist, taxonomist, or applied mathematician) is a judgment call. But the vast majority of cases were clear-cut, and the results don’t change appreciably if the borderline cases are discarded

I recorded each newly hired ecologist’s gender (man or woman), as judged from name, photographs, and pronouns in social media profiles if available. A few new hires also volunteered their gender, unasked for, in emails to me. Using a gender binary, and judging gender from names and photographs, is an imperfect approach, but there was no alternative, and the error rate should be low. I did not attempt to judge race/ethnicity, because that can’t be reliably determined from names and photographs.

I also compiled other various pieces of information about each new hire and the hiring institution. For instance, in the final two years I recorded Carnegie class of US institutions, with Canadian institutions being assigned to the most appropriate Carnegie class when that was clear (e.g., the University of Toronto is the equivalent of an R1 institution). The precise additional information recorded varied from year to year. Some information was compiled only for a subset of new hires because it was time-consuming to compile or could not be determined for all hires.

One year, I read a sample of tenure-track ecology faculty job ads, recording information about the job requirements.

Finally, I compiled information on number of applicants per position from ecoevojobs.net, and by inviting readers of the Dynamic Ecology blog to anonymously share firsthand knowledge. And I compiled information on applicant and search committee behavior by polling readers of the Dynamic Ecology blog on their firsthand knowledge. These polls are not random samples from any well-defined statistical population, but they comprise a usefully-large set of anecdotes from a reasonably diverse set of people.

I present the results as a series of questions and answers. For the answer to each question, I briefly summarize the data source and sample size. For links to the underlying data, deeper analysis, and further discussion, see https://dynamicecology.wordpress.com/ecology-faculty-job-market-data/.

**What fraction of ecology faculty job seekers obtain TT faculty positions?**

*Data: Data on the post-Ph.D. career trajectories of thousands of ecologists, compiled by the US National Science Foundation (NSF), analyzed by Hampton and Labou (2017), and further analyzed by the author*

At least 1/3 of ecology faculty job seekers obtain TT faculty positions; the true fraction may be appreciably higher than that. Hampton and Labou (2017) reported data from the NSF Survey of Doctoral Recipients, a huge stratified random sample of doctoral recipients in all scientific and engineering fields, including ecology. According to Hampton and Labou (2017), 4826 people got US PhDs in ecology from 2000-2011. As of 2013, 10.8% of those 4826 people were postdocs, 3.5% were involuntarily unemployed, and 40.6% of the remainder were in tenured or tenure-track faculty positions. That implies that 1/3 of US ecology Ph.D. recipients from 2000-2011 were in tenured or tenure-track faculty positions as of 2013. The fraction who eventually obtained TT faculty positions will be a bit higher than that, because some of those Ph.D. recipients surely obtained TT faculty positions after 2013.

Further, some unknown fraction of Ph.D. recipients who did not obtain TT faculty positions presumably never wanted or sought a TT faculty position. So the fraction of ecology Ph.D. recipients who want TT faculty positions and go on to obtain them is even higher than 1/3. This may seem surprisingly high, but it is consistent with other sources of information. As discussed later in this paper, self-reported data from ecology faculty job seekers indicates that >42% of them receive at least one TT job offer.

Widely quoted data from other sources indicate that a much smaller fraction of biology Ph.D. recipients go on to TT faculty positions. But data for the field of biology as a whole are not representative of ecology specifically. Many biology Ph.D.s are granted in biomedical and biotechnological fields. Recipients of Ph.D.s in those fields often go on to private sector employment, leaving a smaller fraction to go on to TT faculty positions. Annual rates of Ph.D. production and TT faculty job openings also vary greatly among fields of biology.

The take home message here is not that the ecology faculty job market is uncompetitive. It is competitive: TT ecology faculty job seekers outnumber ecology faculty jobs, and so a substantial fraction of TT ecology faculty job seekers will never obtain a TT faculty position. But the ecology faculty job market is not as competitive as one might think.

**How many ecology faculty job ads welcome or require applicants who collect their own field data? How many welcome or require applicants with quantitative skills?**

*Data: 75 tenure-track ecology faculty job ads listed on ecoevojobs.net as of Sept. 6, 2017*.

Ecology faculty job ads vary widely in their job requirements. Contrary to some anecdotal speculation on social media, most ecology faculty job ads continue to welcome or even require applicants who collect their own field data, and only a small minority require advanced quantitative skills beyond the typical training of North American Ph.D. holders in ecology. 64% of ads welcomed, encouraged, or required applicants who collect their own field data. Only 40% mentioned any quantitative skill, most commonly the ability to teach introductory undergraduate biostatistics. Only 21% of ads mentioned or required “strong” quantitative skills, or specified specific advanced quantitative skills such as Bayesian state space modeling.

**How many TT ecology faculty hires are there at different types of institution?**

*Data: 344 TT assistant professor positions in ecology and allied fields filled in 2016-17 and 2017-18*.

35% of TT ecology hires were at R1 universities (the most research-intensive Carnegie category), 25% were at R2 or R3 universities, 24% were at M1-M3 universities (“master’s” universities lacking substantial Ph.D. programs), 16% were at bachelor’s colleges (colleges lacking substantial graduate programs), and one was at a tribal college. These data reflect variation in the numbers of institutions in different Carnegie categories, and the fact that research universities tend to employ more faculty than other institutions.

**How many applications does the typical TT faculty position in ecology receive?**

*Data: 86 TT positions filled from 2014-15 to 2017-18 (48 from anonymous information on ecoevojobs.net, 38 from readers of Dynamic Ecology)*.

Number of applicants per position varies widely: median 100 applicants, middle 50% 61-175, range 12-1,000 (note that the position that received 1,000 applicants was a broad cluster hire). Positions on the coasts (especially the Pacific Northwest), and at R1 universities, tend to receive more applicants than others. Presumably at least in part for the same reasons why most people live in coastal states, in the urban areas in which most R1 universities are located. Fisheries/wildlife/natural resources positions tend to receive fewer applicants than others, perhaps because Ph.D. holders in those fields often go to work for government agencies, NGOs, or environmental consultancies, leaving fewer to pursue academic careers.

**How many positions does the typical TT ecology faculty job seeker apply to each year? Does it vary by gender?**

*Data: 364 anonymous ecology faculty job seekers on ecoevojobs.net from 2009-10 to 2017-18*.

The typical ecology faculty job seeker applies to 10.5 TT faculty positions annually (median; mean=15.8), but there is wide and highly skewed variation. Many job seekers reported only a few applications annually, but a minority reported many (middle 50% 4-21.8 applications annually, range 1-100).

Men and women ecology faculty job seekers reported submitting similar numbers of applications annually (means of 17.1 for men, 16.8 for women).

**What types of positions do TT ecology faculty job seekers apply to?**

*Data: anonymous reports of 242 ecology faculty job seekers on ecoevojobs.net from 2015-16 to 2017-18*.

81% of ecology faculty job seekers said or implied that they applied to faculty positions at research universities (e.g., by saying that they applied to “everything” or “broadly”), with 55% explicitly saying they’d applied to R1 universities. In contrast, less than half said or implied that they applied for faculty positions at bachelor’s colleges.

These data are consistent with the fact (discussed elsewhere in this piece) that faculty positions at R1 universities tend to receive more applications than positions at other types of institution. However, these data do not necessarily indicate a widespread preference among ecology faculty job seekers for research-intensive careers, for two reasons. First, as noted elsewhere in this piece, there are many more jobs at research universities than at bachelor’s colleges. Second, there are many reasons why research university jobs might attract more applicants. For instance, research universities tend to be concentrated in urban areas where many job seekers might prefer to live for various reasons.

**How do ecology faculty job applicants typically customize their applications? How do ecology faculty search committees expect them to be customized?**

*Data: survey of 102 current or former ecology faculty job seekers, and 35 recent ecology faculty search committee members, on the Dynamic Ecology blog*

50% of ecology faculty job seekers tailored their applications to the hiring institution. 32% tailored their application to the hiring institution only when it was a job they really wanted. 17% tailored their applications to the *type* of institution, but not to individual hiring institutions. Job seekers using all levels of customization reported receiving offers, with those doing less customization reporting slightly more offers on average. I suspect that applicants who do less customization of each application may apply to more positions and so receive more interviews and offers. Applicants who do less customization may also be stronger applicants, and so not need to do as much customization in order to get interviews and offers.

72% of search committee members preferred applications tailored to the hiring institution, with many of those viewing such tailoring as an “honest signal” of the applicant’s seriousness of interest in the position. That preference for applications tailored to the hiring institution was appreciably more common, and was more likely to be seen as “essential”, among search committee members at non-R1 institutions. Perhaps because smaller, less well-known institutions have more reason to worry that applicants will not understand the job requirements or the mission of the institution.

Drilling down, large majorities of ecology search committee members prefer applications that respond to the specifics of the job ad, briefly mention how the applicant would use the institution’s facilities, and describe how the applicant’s teaching and research would be tailored to the institution’s students and mission. The latter is an especially desired form of customization among search committee members at non-R1 institutions. About half of search committee members also want applicants to read the course catalog and identify specific courses they could teach or develop. Applicants most often report using the same forms of customization that search committee members want to see. The exception was that a substantial minority of applicants reported identifying specific faculty members at the hiring institution with whom they’d hope to collaborate, a form of customization that only 17% of search committee members wanted to see.

The take-home messages here for applicants is that it’s common, and worth your while, to tailor your application at least to the type of hiring institution. Tailoring to the individual institution might be warranted if it’s a job you really want or it’s a not a large research university job. But don’t waste time proposing collaborations with specific faculty. And if you feel you need to make a choice between submitting fewer, more heavily customized applications or more, less heavily customized applications, you should probably choose the latter (see below for data demonstrating the importance of submitting many applications).

**How many interviews and offers can the typical TT ecology faculty job seeker expect to receive in a given year? Are there any variables that predict the number of interviews and offers a faculty job seeker will receive?**

*Data: 215 anonymous ecology faculty job seekers on ecoevojobs.net from 2009-10 to 2017-18*.

Ecology faculty job seekers typically report receiving 0-3 phone/skype interviews annually, with those who submitted more applications reporting more phone/skype interviews on average, and women reporting more phone/skype interviews than men on average (Fig. 1). Women reported receiving a mean of 3.0 phone/skype interviews annually, vs. 2.0 for men (medians were 2 and 1, respectively). Among applicants who reported 5 or more phone/skype interviews in a single year, women outnumbered men 30 to 14. The increase in number of interviews with number of applications appears to level off beyond ∼30 applications/year, presumably because applying for more positions than that often means applying for positions for which you are a poor fit.

**Figure 1.**
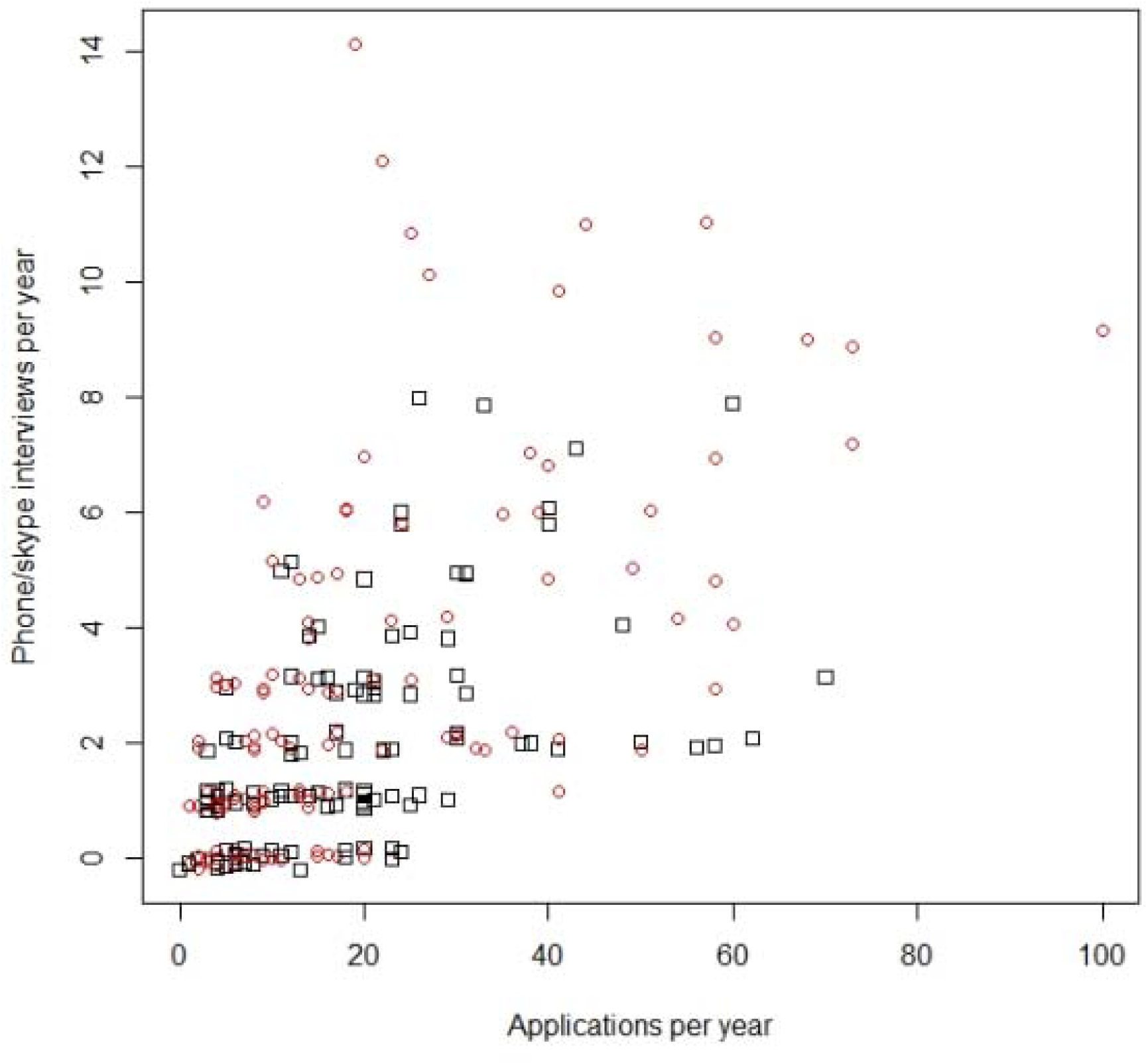
Number of phone/skype interviews received annually by ecology faculty job seekers, as a function of number of applications and gender. Each point gives data for one job seeker in one year. Red circles denote women, black squares denote men. Points are slightly jittered for visibility.

Note that these data likely slightly underestimate the frequency of job seekers receiving no phone/skype interviews. A minority of job seekers did not report the number of interviews they received, and non-reporters tended to be job seekers who applied to few positions and so may have been unlikely to receive any interviews.

Data for number of campus interviews tell a similar story as the data for phone/skype interviews, save that the relationship with number of applications is noisier because campus interviews are rarer than phone/skype interviews (Fig. 2). Women reported receiving an average of 2.4 campus interviews/year (median 2), men reported an average of 1.5 (median 1). 85% of women reported receiving at least one campus interview in a given year, vs. 75% of men. Among applicants who reported receiving at least four campus interviews in a given year, women outnumbered men 26 to 12. The relationship with number of applications again appeared to level off at ∼30 applications. Note that, as with phone/skype interviews, these data may slightly underestimate the frequency of job seekers receiving no campus interviews, due to non-reports.

**Figure 2.**
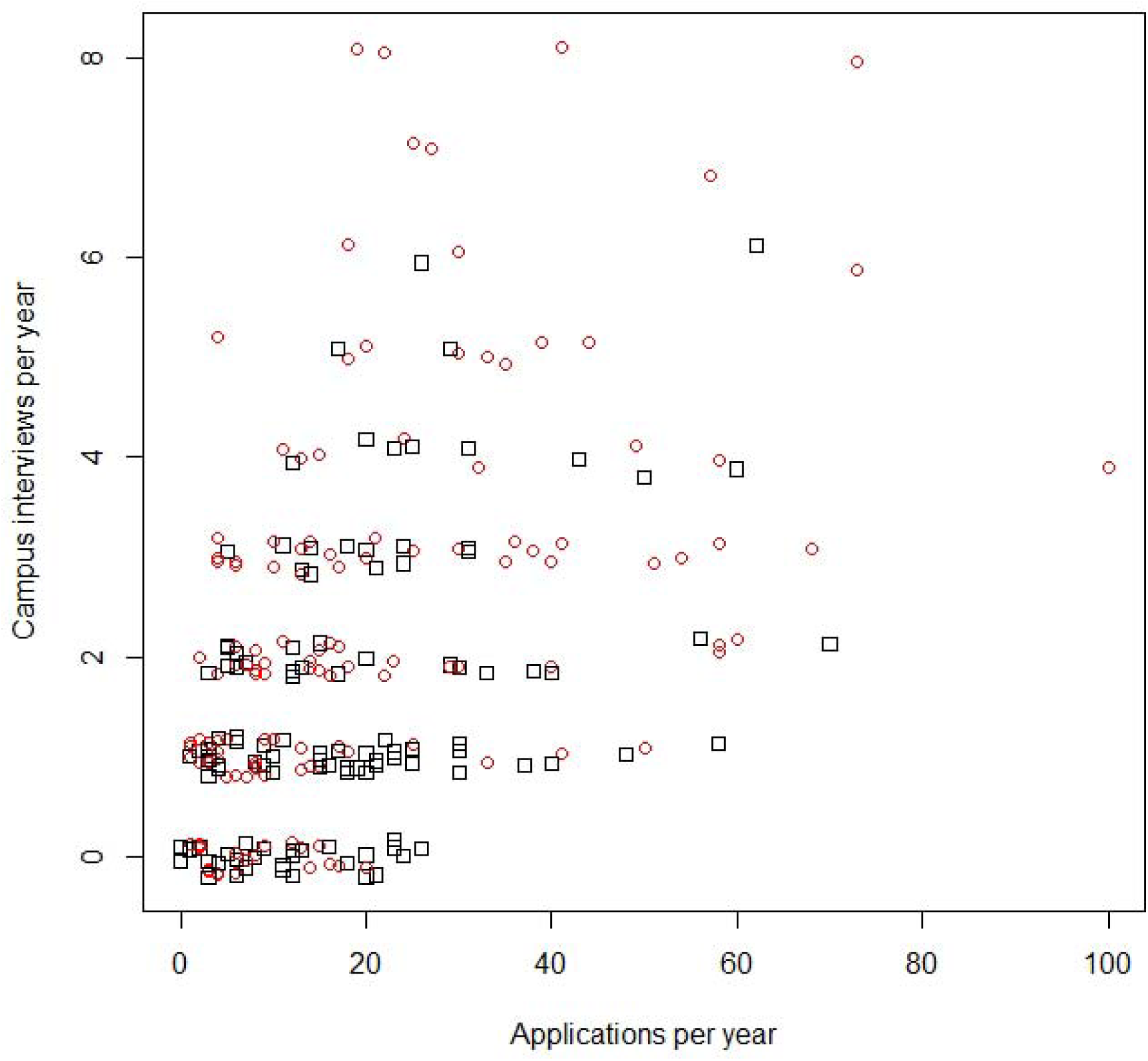
Number of campus interviews received annually by ecology faculty job seekers, as a function of number of applications and gender. Each point gives data for one job seeker in one year. Red circles denote women, black squares denote men. Points are slightly jittered for visibility.

42% of ecology faculty job seekers reported receiving at least one faculty job offer in a given year, and 12% reported receiving multiple offers, counting non-reports of offers as zeroes (Fig. 3). An appreciably higher percentage of women than men reported receiving at least one offer in a given year, and reported receiving multiple offers. The association between number of applications and number of offers is positive but extremely noisy.

**Figure 3.**
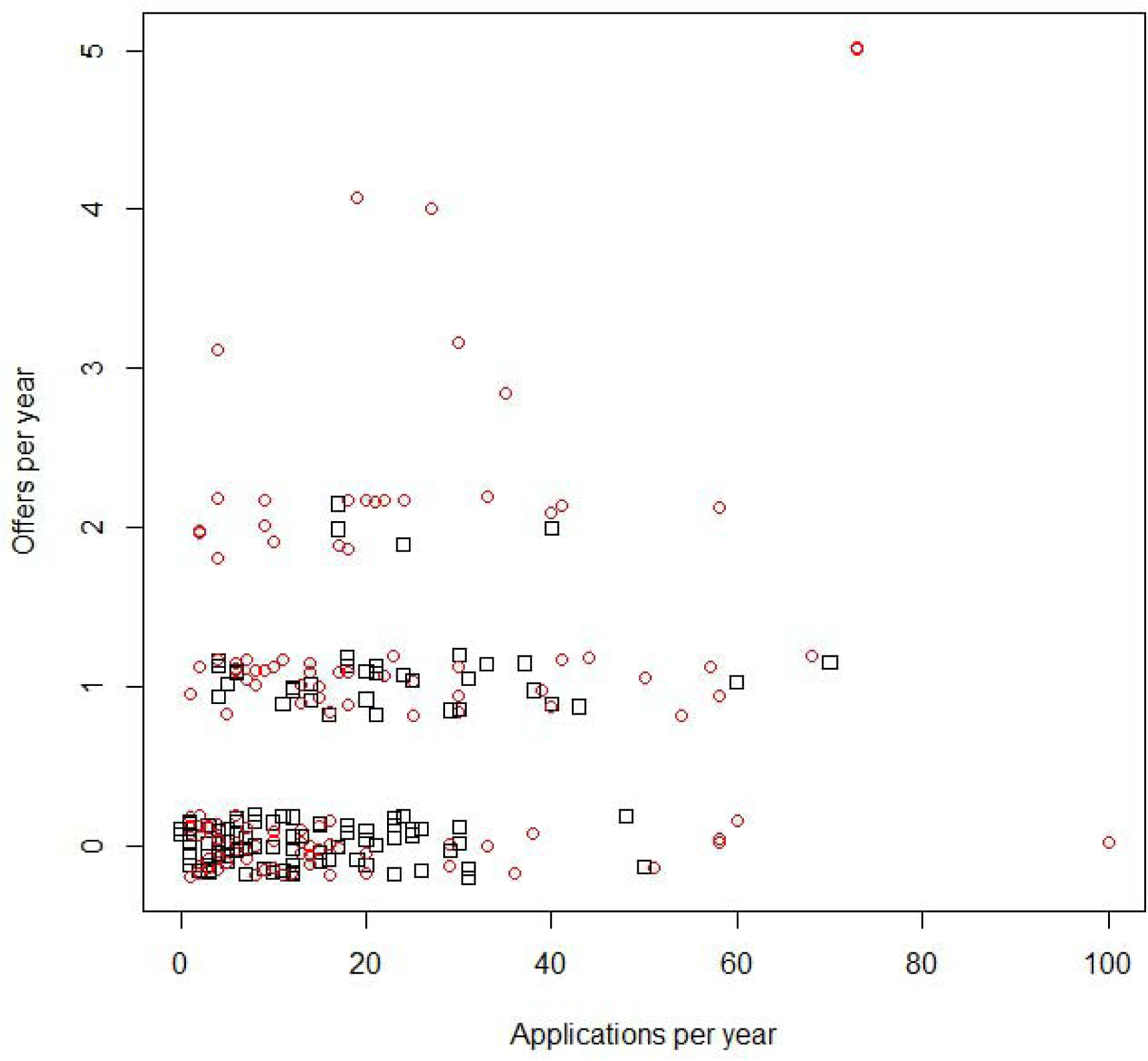
Number of TT offers interviews received annually by ecology faculty job seekers, as a function of number of applications and gender. Each point gives data for one job seeker in one year. Red circles denote women, black squares denote men. Points are slightly jittered for visibility. Non-reports of offers were coded as zeroes.

The results line up with the fact (discussed next) that recently hired TT ecology faculty are 57% women. They reflect the fact that, at a systemic level, most faculty search committees in ecology now take diversity and equity seriously. They want to hire the best people, and the best mix of people, to ensure that their departments and universities are places where all students and faculty can flourish.

No other easily-measurable variables predict the number of interviews or offers ecology faculty job seekers receive, not even if those variables are considered in combination. Variables unrelated to number of interviews or offers include years of post-Ph.D. experience, number of publications, number of first-authored publications, number of major grants received, and number of classes taught.

The take-home lesson for faculty job seekers is that it is to your advantage to apply widely. Although if you find yourself applying for more than ∼30 faculty positions annually, you may wish to consider if some of those applications are long

**What is the gender balance of recently hired TT assistant professors of ecology? Does it vary among types of institution? Is it associated with any measurable gender differences in qualifications? How does it compare to the gender balance of the applicant pool?**

*Data: Gender of 566 TT assistant professors of ecology hired from 2015-16 to 2017-18. Data on institution type and qualifications was collected for those hired in 2016-17 and 2017-18. Also self-reported gender of 401 anonymous ecology faculty job seekers on ecoevojobs.net from 2009-10 to 2017-18*.

Recently hired TT assistant professors of ecology are 57% women. The 566 new hires on which that number is based comprise a large sample of all TT assistant professors of ecology hired in that period, but they’re not a census. If we assume, plausibly, that 750 TT assistant professors of ecology were hired in N. America during that time, then the two-tailed 95% confidence interval ranges from 55-59% women (normal approximation to the binomial distribution for a sample of 566 observations from a finite population of 750). The width of this 95% confidence interval does not widen by more than a percentage point in either direction for any other plausible assumption about the true population size. It is possible that 57% might be a slight underestimate because it proved more difficult to identify hires at small, teaching-intensive institutions, and as shown below those institutions are more likely to hire women than are other types of institution. However, this or any other possible sampling bias would have to be implausibly large to shift the 57% estimate by more than a couple of percentage points in either direction. The fact that this sample comprises a large fraction of all recently hired TT ecology assistant professors leaves little scope for sampling biases to alter the results.

Breaking the gender balance down by year, new hires were 54% women in 2016-16, 59% women in 2016-17, and 58% women in 2017-18. This means that the results are not driven by a single unusual year. The modest year-to-year variation in the gender balance of ecology faculty hires reflects the fact that I was able to identify an average of almost 190 new hires per year, a reasonably large sample. It also reflects the fact that the factors shaping the gender balance of new hires, such as the gender balance of the applicant pool, don’t change much from one year to the next.

The gender balance of recently hired TT ecology faculty covaries with research intensiveness of the hiring institution, although the effect size is modest. Over the three years from 2015-16 to 2017-18, new hires were 54% women at R1 universities or their Canadian equivalents, and 59% women at other types of institution. From 2016-17 to 2017-18 (the only years over which I compiled full Carnegie classification data for all hiring institutions, rather than just R1 vs. non-R1), new hires were 55% women at Ph.D.-granting research universities (i.e. R1, R2, and R3 institutions), with no obvious difference between R1 vs. R2-3, and 63% women elsewhere. These results echo data from other fields. In almost all scholarly fields, women comprise a higher proportion of faculty at more teaching-intensive institutions. They also echo the fact that women comprise a high percentage of school teachers in all OECD countries. Because these trends are not ecology-specific, the reasons for them seem unlikely to be ecology-specific.

Recently hired men and women TT ecology faculty were equally experienced on average. For instance, in 2017-18, the median Ph.D. year for newly hired ecology faculty was 2014 for both men and women, and the means differed by only five months (mean Ph.D. year 2013.4 for men, 2013.8 for women). Recently hired women and men differed a bit on average in some measurable indices of research productivity, such as h-index, but these differences mostly vanished after controlling for the research intensiveness of the hiring institution. Combining data from 2016-17 and 2017-18, men and women hired at bachelor’s colleges had mean Google Scholar h-indices at the time of hiring of 6.9 and 6.6, respectively. At master’s institutions (Carnegie categories M1-M3), the means were 7.7 for men and 6.7 for women. At research universities (Carnegie categories R1-R3), the means were 10.3 for men and 9.4 for women. The remaining small differences in mean h-index have many potential explanations having nothing to do with how well-qualified recent hires are.

Recent TT ecology faculty hires are 57% women even though the applicant pool is a bit short of gender parity. US ecology postdocs were 46% women as of 2013 (Hampton and Labou 2017). And anonymous ecology faculty job seekers on ecoevojobs.net were 47% women over the last decade.

My interpretation of these data is that these days in ecology, multiple excellent women and men apply for most N. American TT faculty positions (a state of affairs that of course reflects everything that shapes people’s career choices and outcomes at the pre-faculty stages). So search committees for most ecology faculty positions will have the opportunity to choose between strong men and women candidates. To help them choose well, search committees these days get trained about bias, diversity, and equity, and have to obey HR rules designed to ensure fairness (e.g., rules obliging search committees to ask all candidates the same questions in the same order during interviews). And many search committees these days are keen for their departments to become more diverse on various dimensions, including but by no means limited to gender. Which is a good thing for them to want. After all, individual faculty don’t exist in isolation from one another. Departments, and the colleges and universities comprised of them, are institutional wholes that are greater than the sum of their parts. Those institutional wholes are best able to teach and inspire the full range of students who come through their doors, and best able to pursue new knowledge, if they’re comprised of diverse, complementary mixtures of people. So gender and other personal attributes are among the many things that search committees consider when they get down to making difficult judgment calls about whom to hire from among (typically) 3-5 well-qualified candidates, each of whom would be an asset to the hiring department in their own unique way (i.e. there’s often not a single “best” candidate on all dimensions). The net outcome is that, at an aggregate statistical level, the proportion of women among recently-hired TT N. American ecology faculty ends up being modestly higher than the proportion of women in the applicant pool, without any appreciable differences in average on-paper qualifications between men and women recently hired at the same type of institution. This is good news, and represents important systemic progress that ecologists should seek to maintain and build upon in order to create a climate in which all ecologists can flourish.

**Where did recent hired ecology professors get their bachelor’s degrees? Did most get their bachelor’s degrees from “top” institutions? Do bachelor’s colleges tend to hire people with bachelor’s degrees from bachelor’s colleges?**

*Data: 75 TT assistant professors in ecology and allied fields hired in 2017-18. Those 75 people include all 23 identified hires at bachelor’s colleges that year, plus 52 randomly-chosen hires at other types of institution*.

Those 75 recent hires got their bachelor’s degrees from 66 different institutions. Only 6 of 75 got bachelor’s degrees from liberal arts colleges or national universities ranked in the 2019 US News and World Report top 30 (a crude proxy for “top” institutions). 84% got their bachelor’s degrees in the US. The remainder received bachelor’s degrees from one of eight other countries. My interpretation of these results is that there good undergraduate students everywhere, some of whom go on to become academic ecologists.

Contrary to what you may have heard, bachelor’s colleges showed no strong tendency to hire graduates of bachelor’s colleges. Only 7/23 bachelor’s college hires received their bachelor’s degrees from bachelor’s colleges. As discussed below, bachelor’s colleges do indeed want to hire faculty who understand their students and their unique institutional missions—but they don’t do that by hiring graduates of bachelor’s colleges.

**Where did recently hired ecology professors get their Ph.D.s? Did most get their Ph.D.s from “top” institutions? Do “top” institutions tend to hire people with Ph.D.s from “top” institutions?**

*Data: 321 TT assistant professors in ecology and allied fields hired in 2016-17 or 2017-18*.

Recently hired ecology professors got their Ph.D.s from a wide range of places: 147 different institutions. No single institution granted more than 3% of the Ph.D.s held by recently hired TT N. American assistant professors of ecology. The diversity of institutions from which recently hired TT ecology faculty received their Ph.D.s is comparable to the tropical tree species diversity on Barro Colorado Island in Panama, as measured by Simpson’s index. This makes ecology very different from the fields of computer science, business, and history, in which the top 10 US Ph.D. programs train >70% of all US TT faculty (Clauset et al. 2015).

86% of recent ecology faculty hires received Ph.D.s from US institutions, 8% from Canadian institutions, the rest received their Ph.D.s from one of ten other countries. Presumably, the rarity of hires with Ph.D.s from outside North America primarily reflects the rarity of applicants from outside North America.

In contrast to the fields of computer science, business, and history (Clauset et al. 2015), highly ranked ecology programs show no obvious tendency to only hire graduates of other highly ranked programs. This is good news, and presumably reflects the fact that faculty search committees have many lines of evidence with which to evaluate applicants—cv’s, publications, research and teaching statements, reference letters, etc. They don’t need to rely on graduate program reputation as a crude proxy to evaluate applicants.

**Do you have to have a paper in Science, Nature, or PNAS to be hired as a TT ecologist? Does the answer depend on whether you’re seeking a position at an R1 university?**

*Data: 314 TT assistant professors of ecology hired in 2016-17 or 2017-18*.

Only 25% of recently hired TT ecology professors had papers in Science, Nature, or PNAS at the time they were hired, and only 12% had first-authored papers in those journals. Even at the most research-intensive institutions, papers in those journals are far from essential to be hired into a TT ecology faculty position. Among recent hires at R1 institutions or their Canadian equivalents, 45% had Science, Nature, or PNAS papers at the time of hiring, and 27% had first-authored papers in those journals.

**Do you have to have a famous Ph.D. supervisor to be hired as a TT ecologist at a research-intensive institution?**

*Data: Attributes of the Ph.D. supervisors of 39 TT assistant professors of ecology hired at R1 universities in 2017-18, and 20 TT assistant professors of ecology hired at bachelor’s colleges in 2017-18*.

Online discussions of the ecology faculty job market occasionally suggest that having had a famous Ph.D. supervisor is common or even essential among recent faculty hires at R1 universities. This is incorrect, for any plausible operational definition of “famous”. The Google Scholar h-indices of the Ph.D. supervisors of recently hired TT ecology faculty at R1 universities vary widely (median 47, range 7-153), do not differ appreciably from the h-indices of the supervisors of recent hires at bachelor’s colleges (median 44, range 29-77), and resemble the h-indices of a representative sample of tenured R1 university ecology professors. That is, these data look more or less as would be expected if recently-hired ecology faculty at all types of institutions had done their Ph.D.s with a random sample of tenured ecology faculty at research universities.

Other operational definitions of “famous” lead to the same conclusion. For instance, only 7% of R1 hires, and no bachelor’s college hires, were supervised or co-supervised by a member of the US National Academy of Sciences.

**How many years of post-Ph.D. experience did recently hired TT ecology professors typically have? How does that compare to the experience level of the applicant pool?**

*Data: 306 TT assistant professors of ecology hired in 2016-17 or 2017-18. For purposes of calculating years post-Ph.D., those hired in 2016-17 were coded as 2017 hires; those hired in 2017-18 were coded as 2018 hires. Also 441 anonymous ecology faculty job seekers on ecoevojobs.net from 2009-10 to 2017-18*.

Recently hired TT assistant professors of ecology typically were about 4 years post-Ph.D. at the time of hiring (mean 4.2 years, median 4). 69% had anywhere from 2-6 years of post-Ph.D. experience, with a range from 0-11 years of post-Ph.D. experience. The majority of ecology faculty job seekers also are 2-6 years post-Ph.D.

These data are consistent with a job market in which most hiring institutions prefer applicants with at least a year or two of post-Ph.D. experience. But once you have a few years of post-Ph.D. experience, the marginal value of additional post-Ph.D. experience appears to be low at best.

**How were recently hired TT ecology professors employed at the time of hiring?**

*Data: 144 TT assistant professors of ecology hired in 2017-18*.

65% of recently hired ecology faculty were postdocs at the time of hiring. 13% were TT or (much more rarely) tenured professors at another institution. 8% were in a non-tenured research-focused position such as “research professor” or “scientist”. 4% were in a non-tenured teaching-focused position such as “teaching fellow” or “instructor”. 3% were visiting assistant professors. The remainder were in other employment, though two of those were also adjunct professors.

One take-home message here for faculty job seekers is that TT faculty positions rarely are filled by people in non-academic employment. You can’t leave academia and expect to return later. Another take-home message for faculty job seekers is not to worry too much about competition from candidates who already hold tenured or TT positions elsewhere. In my experience, applicants who already hold tenured or TT positions comprise only a small fraction of all applicants, are judged according to higher standards than more junior applicants, are more expensive to hire, and are less likely to accept any offer they might receive. For all these reasons, the TT faculty job market in ecology is not dominated by existing TT faculty playing “musical chairs.”

Another take-home message for faculty job seekers is that it’s actually not common for TT faculty to have previously served as visiting assistant professors or in some other exclusively teaching-focused non-TT position. As a faculty job seeker, don’t assume that you “have” to take a full-time non-TT teaching position in order to eventually land a TT faculty position, not even a TT position at a teaching-intensive institution. Teaching-intensive institutions do indeed expect their TT faculty to have substantial teaching experience (see below). But most recently hired ecologists acquired that experience in some other way besides taking full-time non-TT teaching positions.

**What teaching experience did recently hired TT assistant professors of ecology have?**

*Data: 29 TT assistant professors of ecology hired at R1 universities in 2017-18, and 16 TT assistant professors of ecology hired at bachelor’s colleges in 2017-18*.

Almost all recently-hired TT ecology professors had some form of teaching experience and/or pedagogical training. Only two R1 university hires listed no teaching experience or pedagogical training on their cv’s. However, I suspect that even they likely served as teaching assistants. 23/29 R1 hires and 12/16 bachelor’s college hires listed teaching assistant experience on their curriculum vitae, and I suspect that most or all of the others had it but didn’t bother to list it.

Experience as an instructor of record is almost essential to be hired for a TT faculty position at the most teaching-intensive institutions. 14 of 16 recent hires at bachelor’s colleges served as instructors of record before being hired, often for multiple courses. Of the other two, one served as a co-instructor. The other designed and taught a course for high school students, an unusual but substantial form of teaching experience. Experience as an instructor of record was rarer but far from unheard of among recent R1 university hires: 12/29 had it.

Guest lecturing is a common form of teaching experience among R1 hires; 16/29 listed guest lecturing on their cv’s. Only 4/16 bachelor’s college hires did so, perhaps because experience as an instructor of record makes guest lecturing experience redundant.

Guest lecturing and teaching assistant experience are the most common forms of teaching experience among R1 hires. But only 9/29 R1 hires had teaching experience limited to teaching assistantships and/or guest lecturing.

Only 3/16 bachelor’s college hires and 4/29 R1 hires listed any formal pedagogical training on their cv’s.

Based on these data, I would strongly advise faculty job seekers to obtain experience as an instructor of record if they plan to seek a teaching-intensive faculty position, and not to bother applying for teaching-intensive faculty positions unless they have that experience or equivalent experience. However, faculty job seekers seeking TT faculty positions at research universities should carefully consider whether serving as an instructor of record would take too much time away from research, relative to the benefit to their pedagogical skills and job prospects. Attending pedagogical training workshops and short courses might be a less time-intensive way to acquire some teaching skills and signal to future employers that you take teaching seriously.

**Are many TT ecology faculty positions filled by “inside” candidates?**

*Data: 35-326 TT assistant professors of ecology hired in 2016-17 and 2017-18*.

A common worry among faculty job seekers who comment on ecoevojobs.net is that they will lose out to “inside” candidates whose current or past connections to the hiring department will work in their favor, even if they are not the best candidates. Faculty job seekers who have this worry may wish to consider that they’re worrying about a very unlikely possibility. Very few TT assistant professor positions in ecology and allied fields are filled by a candidate with any current or previous connection to the hiring department:

- 3% of new hires got a bachelor’s degree from the hiring institution (sample size: 77 new hires in 2017-18)
- 0.3% of new hires got a Ph.D. from the hiring institution (sample size: 307 new hires in 2016-17 and 2017-18)
- 4% of new hires were employed by the hiring institution at the time of hiring (sample size: 326 new hires in 2016-17 and 2017-18)
- No new hires were employed by the hiring institution sometime after obtaining their Ph.D.s, but not employed by the hiring institution at the time of hiring (sample size: ∼150 new hires in 2017-18)
- No new hires lacking a previous educational or employment connection to the hiring institution had ever co-authored a paper with someone in the hiring department (sample size: 35 new hires in 2017-18).

These data indicate that faculty job seekers rarely obtain a TT faculty position by first developing an educational or professional connection with the hiring department. This is consistent with my own experience. My own experience is that it is extremely rare for TT faculty positions to be intended for a specific “inside” candidate from the get-go. For other faculty positions, “inside” candidates are a small fraction of all applicants at most, and are evaluated on a level playing field with the other applicants.

**What is the typical Google Scholar h-index of recently hired TT assistant professors of ecology?**

*Data: Google Scholar h-indices of 264 TT assistant professors of ecology hired in 2016-17 or 2017-18*.

The Google Scholar h-index is a crude summary measure of publication productivity and the frequency with which those publications are cited. It has numerous flaws and limitations, for instance weighting co-authored publications the same as sole- or first-authored publications. And it is not used by ecology faculty search committees to evaluate applicants, as far as I am aware. But it may be loosely correlated with attributes that search committees do consider, and it might be useful to prospective faculty job seekers to know if their own h-indices fall within the typical range for new hires. Looking at h-index data also is a convenient way to test the widespread impression that you have to have many publications in order to be competitive for a TT faculty position in ecology. Your h-index cannot exceed your number of publications.

Recently hired TT assistant professors of ecology typically have an h-index of 8 (median; mean=8.7), but the values range widely. The middle 50% of the distribution ranges from 6-11, and the full range is 1-24. Some of this variation reflects the fact that h-indices run higher on average among new hires at Ph.D.-granting research universities compared to new hires at other institutions (though the distributions do overlap). But h-indices often vary by a factor of two even among new hires of similar experience levels who were hired into the same department.

Note that the wide spread of h-indices does not primarily reflect variation among recent hires in years of post-Ph.D. experience. H-index is only loosely positively correlated with years of post-Ph.D. experience among these 264 recent hires.

Note as well that not all new hires have Google Scholar pages, and those who don’t are concentrated at less research-intensive institutions. For that reason, these data likely slightly overestimate the typical h-index of recently hired TT assistant professors of ecology.

**How many first-authored publications do you need to have to be hired as a TT ecologist at an R1 university? How many of those first-authored publications need to be in “leading” journals? How do the first-authored publication counts of recent hires at R1 universities compare to those of the applicant pool? Do first-authored publication counts of applicants vary by gender?**

*Data: First-authored peer-reviewed paper counts of 39 TT assistant professors of ecology hired at R1 universities or their Canadian equivalents in 2017-18. “Leading” journals were operationally defined as those with a 2-year impact factor* ≥*3, regardless of the journal’s field. Book chapters, government technical reports, book reviews, letters to the editor, comments, etc., were not counted. Also first-authored publication counts of 442 anonymous ecology faculty job seekers on ecoevojobs.net from 2009-10 to 2017-18*.

Ecology faculty search committees at research universities don’t base their hiring decisions on crude quantitative metrics like publication counts. But they do want to hire faculty who will lead high quality, high impact research programs. One line of evidence for the ability to do that is the applicant’s record of first-authored publications. Publication count data are time-consuming to compile, so I compiled them only for recent hires at R1 universities. These counts should provide an upper bound on the publication counts of ecologists hired at less research-intensive institutions.

Recently hired TT assistant professors of ecology at R1 universities varied widely in how many first-authored papers they’d published at the time of hiring, and how many of those first-authored papers were in “leading” journals. The typical recent R1 hire had 7 first-authored papers (median; mean=9.5), with a wide spread around the typical value (middle 50% =5-11, range=3-38). The typical recent R1 hire had 4 first-authored papers in leading journals (median; mean 4.4), again with a wide spread around the typical value (middle 50% 2-6, range 0-14). No, that last range is not a typographical error. There were actually multiple ecology faculty hired into TT assistant professor positions at R1 universities in 2017-18 with no first-authored papers in journals with two-year impact factors ≥3.

Note that this wide spread does not primarily reflect variation among recent R1 hires in years of post-Ph.D. experience. Counts of first-authored publications, and first-authored publications in leading journals, were only loosely positively correlated with years of post-Ph.D. experience among these 39 recent R1 hires.

Part of the explanation for why some recent hires had few or no first-authored papers in “leading” journals is because some work in specialized subfields in which there are few or no journals with two-year impact factors ≥3 (e.g., ecological entomology, fisheries). However, that is only part of the explanation; some recent hires with <2 first-authored publications in “leading” journals were hired into faculty positions in “ecology” or broad subfields.

The first-authored publication counts of ecology faculty job seekers as a group are similar to those of recently hired TT ecology faculty at R1 universities (median 7, mean 8.6, middle 50% 5-10, range 0-50). Women ecology faculty job seekers have fewer first-authored publications than men on average (mean 6.8 for women, 10.5 for men; medians 6 and 9, respectively).

The take-home message here for faculty job seekers is not that first-authored publications and publication venue don’t matter for faculty hiring decisions at research universities, but that many other factors matter as well.

## Concluding advice for ecology faculty job seekers

### Variability is the rule

Recently hired ecology faculty vary a lot on every easily-measurable dimension. The range of qualifications that makes an applicant “competitive” for any given position is wider than many ecology faculty job seekers—and even many ecology faculty—probably realize. So if there’s a faculty position you think you might want, go ahead and apply for it. Don’t take yourself out of the running by assuming that you’re not “competitive”.

### Fit matters

Most easily-measurable attributes of faculty job seekers have no predictive power for how many interviews or offers they will receive. That’s because faculty search committees don’t decide who to interview or hire based on crude quantitative metrics like years of experience, publication count, h-index, etc. Rather, they evaluate applicants holistically, paying great attention to attributes that can’t be measured, particularly fit to the position. This doesn’t mean hiring decisions are arbitrary, of course; some applicants really do fit any given position better than others.

### Don’t be more pessimistic than the data warrant

The ecology faculty job market is competitive, and being on the market can be very stressful. But in polls conducted on Dynamic Ecology, I’ve noticed a consistent pattern: ecologists are consistently more pessimistic about the ecology faculty job market than is warranted by the data. As a group, ecologists seriously underestimate the percentage of women among recently hired ecology faculty (the modal guess is 40% women). They vastly overestimate the minimum number of first-authored publications required to be hired as an ecology professor at an R1 university. Substantial pluralities think that most recently hired ecology faculty got their Ph.D.s from a relatively small number of “elite” institutions, and that “elite” institutions only hire ecology faculty with Ph.D.s from “elite” institutions. And most overestimate the fraction of ecology faculty positions filled by current employees of the hiring institutions. This misdirected pessimism seems unfortunate. Data can’t tell anyone how to feel, and there absolutely are good reasons why anyone on the ecology faculty job market might feel stressed. But hopefully these data will reassure some job seekers and help them avoid baseless sources of stress.

### All faculty job seekers are in the same boat

In a competitive faculty job market, it’s easy to fall into the trap of believing that people like you—people with your interests, skills, experience, and connections—aren’t getting many or even any offers. But here’s the thing: everybody thinks that—and everybody is right. On a competitive job market, everybody has it tough. Women. Men. Quantitative ecologists. Field ecologists. People with connections to the hiring department. People with no connections to the hiring department. More experienced people. Less experienced people. Etc. So try not to resent faculty job seekers who differ from you in some way; they’re all in the same, competitive boat as you.

## Footnotes

1 Throughout, I restrict attention to North America. These results should not be assumed to generalize to other places.

## Notes

https://dynamicecology.wordpress.com/ecology-faculty-job-market-data/

